# Raw wastewater irrigation for urban agriculture in Africa increases the diversity of transferable antibiotic resistances genes in soil, including those encoding ESBLs

**DOI:** 10.1101/615922

**Authors:** B. P. Bougnom, S. Thiele-Bruhn, V. Ricci, C. Zongo, L.J.V Piddock

## Abstract

A metagenomic study was conducted to investigate the impact of raw wastewater use for irrigation in urban agriculture on the development of bacterial resistance in soil. Soil samples were collected in two African countries, from three different cities (each with irrigated and non-irrigated plots). Basic physical and chemical analysis were conducted, and the presence of selected antibiotic residues was assessed. Microbial DNA was extracted, quantified and sequenced. Microbial population structure and function, presence of horizontally transferable antibiotic resistance genes and *Enterobacteriaceae* plasmids replicons were analysed using bioinformatics. The relative prevalence of *Proteobacteria* and *Bacteroidetes* and sequence reads coding for microbial adaptation and growth were higher in irrigated fields; 33 and 26 transferable ARGs were found in irrigated and non-irrigated fields sequence reads, respectively. Extended spectrum β-lactam genes identified in irrigated fields included *^bla^*CARB-3, *^bla^*OXA-347, *^bla^*OXA-5 and *^bla^*Rm3. Concentration of sulfamethoxazole, ciprofloxacin and enrofloxacin in soils influenced the selection of antibiotic resistance genes encoding resistance against amphenicol, β-lactams, and tetracyclines. Ten *Enterobacteriaceae* plasmid amplicon groups were identified in the fields, five were common to both, two (IncW and IncP1) and three (IncY, IncFIB and IncFIA) were found in irrigated and non-irrigated fields, respectively.

In conclusion, wastewater irrigation affected both soil microbial diversity and functions. Irrigated fields have more diverse transferable antibiotic resistance genes, including ESBL genes that encode resistance to β-lactams antibiotics, except cephamycins and carbapenems. Even more, critical concentrations of antibiotic residues select for multiple and cross resistance. The findings from African cities show that wastewater irrigation in urban agriculture presents a serious public health risk for farmworkers and consumers by spread of bacterial resistance.

## 1. Introduction

Antibiotic-resistant bacteria (ARB) are a serious health problem even at the world’s most advanced medical centres (Piddock, 2012). In 2011 an epidemic of *Escherichia coli* infections caused by contaminated bean sprouts affected up to 5,000 people in Europe, with over 48 deaths (Buchholz et al., 2011). These two examples emphasize that morbidity, mortality, and the associated economic burden owing to bacterial resistance which cost $20 billion in health care costs annually in the US, are likely to vastly increase during the next decade. To efficiently tackle the increasing bacterial resistance, environmental, agricultural, and medical aspects need to be handled at a global scale (Wellington et al. 2013).

Arable lands reported to be irrigated with wastewaters worldwide cover approximately 20 million hectares; which equates to 10% of the total global irrigated land (Mateo-Sagasta et al. 2011). Wastewaters can contain high concentrations of antibiotics originating from slaughterhouses, private use and applications in hospitals along with numerous bacterial species, some of which are pathogenic or resistant to antibiotic agents or both, and thus may have a pronounced effect in that sense on soil microbial communities subsequent to soil irrigation with wastewater (Dickin et al. 2016). Whereas in the past, several research teams have addressed questions related to the application of manure which is contaminated with antibiotics used in husbandry, much less is known on the effect of raw wastewater irrigation on the development of ARB in irrigated fields, although raw wastewater irrigation is often applied in low and middle income countries (LMIC) as a cheap alternative in case of water scarcity and to costly commercial fertilizers. Wastewater irrigation in urban agriculture has been a common practice for decades in many cities in LMICs (Adegoke et al. 2018).

The release of pharmaceuticals in the environment selects for drug resistant bacteria (Andersson and Hughes, 2010). To combat antibiotics in ecosystems, bacteria have evolved a plethora of different antibiotic resistance genes (ARGs) of which many are mobile and can easily spread between species including human and animal pathogens. Resistant telluric bacteria can transfer ARGs to pathogenic bacteria by horizontal genes transfer (Forsberg et al. 2012). There is a crucial need to identify the principal reservoirs of ARGs in humans, animals and environment, since there is insufficient information about the conditions and factors that lead to the mobilization, selection and movement of resistant drug bacteria into and between environment, human and animal populations (Wellington et al. 2013).

Metagenomics allow to understand the factors driving ARGs and the different antibiotic resistance mechanisms (Amos et al. 2014). The use of a metagenomic approach gives us a realistic opportunity to alternative strategies for prevention against present and future clinically relevant cases. It will also aid in the design of inhibitors preventing ARGs transfer between bacteria, discovery and production of new compounds less susceptible to existing resistance mechanisms (Alves et al. 2018).

In this study, we assess the impact of raw wastewater use in urban agriculture on microbial communities and development of bacterial drug resistance in agricultural fields. It is postulated that in agricultural field irrigated with raw wastewater, there are bacteria adapted to survive antibiotics exposure by vertical and horizontal gene transfer, with a high number containing clinically relevant ARGs.

## 2. Material and methods

### 2.1. Experimental design and soil sampling

The experiment was conducted in three cities, in two African countries, namely Ouagadougou (46°38′ N, 11°29′) in Burkina Faso, Ngaoundere (46°38′ N, 11°29′) and Yaounde (46°38′ N, 11°29′) in Cameroon (Figure S1). Their respective annual mean of temperature and precipitations are for: Ouagadougou (30°C; 867 mm); Ngaoundere (22°C; 1497mm) and Yaounde (24°C; 1628 mm), respectively. At each city two blocks were investigated, comprising three agricultural fields that were irrigated (IRI) with raw wastewater, and as control soils, 500 m away, three non-irrigated agricultural fields (NIR), with comparable soil properties. We had Ouagadougou (IRI1 and NIR1); Ngaoundere (IRI2 and NIR2), and Yaounde (IRI3 and NIR3). The agricultural fields were approximately 0.2 ha each and watered manually with watering cans. In each field, 100 g of soil were sampled from 0-20 cm depth, using soil cores. This was repeated at 10 randomly distributed places within each field. Replicate samples were pooled together, receiving 1 kg-composite samples. The samples were transported on ice and stored at −80°C until further analysis.

### 2.1. Soil physical and chemical analysis

Soil pH was measured in a 1:2.5 (soil: demineralised water) ratio using a glass electrode. Total C and N were analysed using a TOC-VCPN-analyzer (Shimadzu, Duisburg, Germany).

Soil antibiotic residues were determined according to Blackwell et al. (2004). Briefly, 4 g of air-dried soil were weighed into 10 ml centrifuge tubes, and 5 ml of extraction buffer (0.1 M McIlvaine buffer (Na2HPO4 and citric acid at pH 7)/0.1 M EDTA/MeOH 25:25:50 v/v) were added. The tubes were vortexed for 30 s and placed in an ultrasonic bath for 10 min, and then centrifuged at 1160 g for 15 min. The supernatant was collected, and the operation was repeated twice. The collected supernatants were pooled together and diluted to 400 mL with distilled water and acidified to pH 2.9 with phosphoric acid prior to solid phase extraction (SPE). SAX and HLB SPE cartridges (Thermofisher, Massachusetts, USA) were set up in tandem for SPE. The cartridges were conditioned with 5ml methanol then conditioned with 5ml buffer (2.5 ml 0.1M NaOAc, 5ml distilled water and 2 ml 20% methanol) at a rate of 2 ml min^-1^. Thereafter, the supernatants were loaded at a flow rate of 10 ml min^-1^. Then, the SAX cartridges were removed and the HLB cartridges were washed with 5ml conditioning buffer, Thereafter, the HLB cartridges were air dried for 10 min and antibiotic residues were eluted with 2 × 1 ml of methanol. The eluates were evaporated using a rotavapor rotor, introduced in 1 ml vial tubes for further analysis. Antibiotic concentrations in the extracts were determined according to Michelini et al. (2012), using a Shimadzu LC-20 HPLC (Shimadzu, Duisburg, Germany) coupled to an API 3200 LC–ESI-MS/MS (Applied Biosystems/MDS Sciex Instruments, Toronto, Canada). The analysed antibiotic compounds included sulfadimidine, sulfadiazine, sulfamethoxazole, ciprofloxacin, enrofloxacin, chlortetracycline, oxytetracycline, tetracycline, trimethoprim and tylosin.

### 2.2. Microbiological analysis

#### 2.2.1. Soil Biomass Purification

To collect mainly the bacterial cells from the different soils, soil biomass purification was conducted according to Sentchilo et al. (2013). Briefly, 15g soil samples were homogenized by magnetic stirring for 15min, in ice-cold poly (beta-amino) esters (PBAE) buffer (PBAE buffer is 10mM Na-phosphate, 10mM ascorbate, 5mM EDTA, pH 7.0), at 10 ml g^-1^ of soil. Low speed centrifugation in 50-ml conical tubes at 160 g for 6 min was used to remove coarse particles, big eukaryotic cells and bacterial flocks. The collected supernatants were centrifuged at 10000 g for 5 min to pellet the microbial biomass for further analysis.

#### 2.2.2. DNA extraction and quantification, and high-throughput sequencing

Soil DNA was extracted using the DNeasy PowerSoil Kit (Qiagen, Germany) according to the manufacturer’s instructions. DNA concentration was determined by using the Quant-iT PicoGreen dsDNA Assay Kit, and the Qubit™ 3.0 Fluorometer (Qubit, Life Technologies, USA)The three DNA samples extracted from each block were pooled together in equal nanogram quantities. Six DNA samples representative of the three cities were sent to Edinburgh Genomics for high-throughput sequencing. Sequencing was conducted using Illumina Hiseq4000 (Illumina, Inc, USA), TruSeq DNA Nano gel free library (350 bp insert) was used to prepare the libraries. Raw data consisted of 190.5 Gb sequences. The metagenomic data have been deposited at the National Center for Biotechnology Information (NCBI), Sequence Read Archive (SRA) under project accession number PRJNA358310.

#### 2.2.3. Taxonomic and functional annotations

The raw metagenomic sequences were uploaded to the metagenomics RAST server (MG-RAST) version 4.0.3 (Glass et al., 2010). Microbial community profiling was conducted using the SEED database, and metabolic profiles assignments were annotated against SEED subsystems database. The microbial functional profiles were annotated against SEED subsystems (collections of functionally related protein families). Both microbial and metabolic profiles were generated using E-value cut-off 10^-5^, at a minimum identity of 80%, and a minimum alignment length of 20 amino acids (Glass et al., 2010).

#### 2.2.4. Identification and quantification of antibiotic resistance gene markers

Short Better Representative Extract Dataset (ShortBRED, Kaminski et al., 2015) was used to identify and quantify antibiotic resistance genes (ARGs) from the metagenome. ShortBRED profiles protein family abundance in metagenomes in two-steps: (i) *ShortBRED-Identify* isolates representative peptide sequences (markers) for the protein families, and (ii) *ShortBRED-Quantify* maps metagenomic reads against these markers to determine the relative abundance of their corresponding families based on reads per kilobase million (RPKM). Fragment length ≥ 30 amino acids and ≥ 95% identity was considered positive. The Comprehensive Antibiotic Resistance Database (CARD, McArthur et al., 2013) was used to generate ARG markers, using UniRef50 as a reference protein database. Antibiotic resistance ontology (ARO) numbers in CARD was used to aggregate, annotated and associate the ARGs to the corresponding resistance family.

#### 2.2.5. Identification of plasmid amplicons of clinical relevance

*Enterobacteriaceae* plasmid replicon sequences were downloaded from the PlasmidFinder database 1.3 (https://cge.cbs.dtu.dk/services/PlasmidFinder). The nucleotides sequences were aligned against the metagenomic reads using BLAT. The parameters were BLAT hit with a sequence identity ≥ 80% and E-value cut-off of 10^-5^ (Carattoli et al., 2014).

### 2.3. Data Analysis

The relative abundance of the different bacterial phyla and families of interest, the functional categories present in the metagenomic reads, ARGs, and *Enterobacteriaceae* plasmid replicon groups detected in the raw wastewater samples in each town were compared using the Student’s *t*-test, results were statistically significant at *P* <0.05.

## 3. Results

### 3.1. Antibiotic residues in soil

The values of soil pH and organic C of the soil samples are reported in Table 1. These parameters were slightly higher in irrigated fields, but the difference was not significant between the two farming systems. Concentrations of sulfamethoxazole, enrofloxacin and oxytetracycline were significantly higher in irrigated fields. Enrofloxacin (1.10 ng. g^-1^) and sulfadimidine (0.81 ng. g^-1^) were the most prevalent antibiotic compounds in irrigated and non-irrigated fields, respectively. Soil contents of the other investigated antibiotics ranged from 0.09 to 0.92 and 0.04 to 0.44 ng g-1 in irrigated and non-irrigated fields, respectively (Table 1). Sulfadiazine, chlortetracycline, tetracycline and tylosin residues, however, were not detected in soils of both farming systems. Oxytetracycline was not discovered in non-irrigated fields.

**Table 1.**
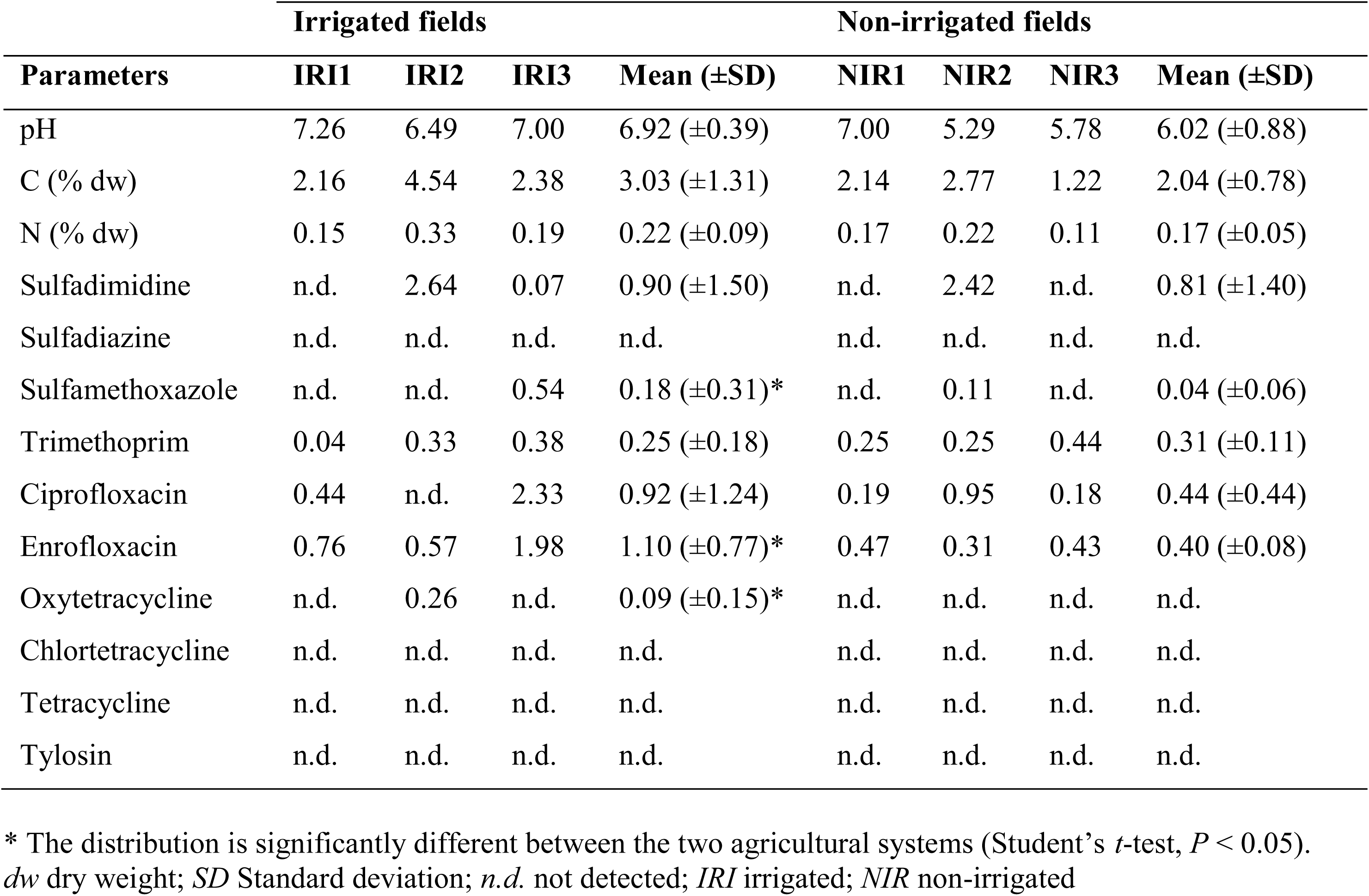
pH, carbon, nitrogen content and concentration of antibiotic residues (ng. g^-1^) in the samples collected in irrigated and non-irrigated fields

| Parameters | Irrigated fields |  |  |  | Non-irrigated fields |  |  |  |
| --- | --- | --- | --- | --- | --- | --- | --- | --- |
| | IRI1 | IRI2 | IRI3 | Mean (±SD) | NIR1 | NIR2 | NIR3 | Mean (±SD) |
| pH | 7.26 | 6.49 | 7.00 | 6.92 (±0.39) | 7.00 | 5.29 | 5.78 | 6.02 (±0.88) |
| C (% dw) | 2.16 | 4.54 | 2.38 | 3.03 (±1.31) | 2.14 | 2.77 | 1.22 | 2.04 (±0.78) |
| N (% dw) | 0.15 | 0.33 | 0.19 | 0.22 (±0.09) | 0.17 | 0.22 | 0.11 | 0.17 (±0.05) |
| Sulfadimidine | n.d. | 2.64 | 0.07 | 0.90 (±1.50) | n.d. | 2.42 | n.d. | 0.81 (±1.40) |
| Sulfadiazine | n.d. | n.d. | n.d. | n.d. | n.d. | n.d. | n.d. | n.d. |
| Sulfamethoxazole | n.d. | n.d. | 0.54 | 0.18 (±0.31)* | n.d. | 0.11 | n.d. | 0.04 (±0.06) |
| Trimethoprim | 0.04 | 0.33 | 0.38 | 0.25 (±0.18) | 0.25 | 0.25 | 0.44 | 0.31 (±0.11) |
| Ciprofloxacin | 0.44 | n.d. | 2.33 | 0.92 (±1.24) | 0.19 | 0.95 | 0.18 | 0.44 (±0.44) |
| Enrofloxacin | 0.76 | 0.57 | 1.98 | 1.10 (±0.77)* | 0.47 | 0.31 | 0.43 | 0.40 (±0.08) |
| Oxytetracycline | n.d. | 0.26 | n.d. | 0.09 (±0.15)* | n.d. | n.d. | n.d. | n.d. |
| Chlortetracycline | n.d. | n.d. | n.d. | n.d. | n.d. | n.d. | n.d. | n.d. |
| Tetracycline | n.d. | n.d. | n.d. | n.d. | n.d. | n.d. | n.d. | n.d. |
| Tylosin | n.d. | n.d. | n.d. | n.d. | n.d. | n.d. | n.d. | n.d. |

\* The distribution is significantly different between the two agricultural systems (Student's *t*-test, *P* < 0.05). *dw* dry weight; *SD* Standard deviation; *n.d.* not detected; *IRI* irrigated; *NIR* non-irrigated

### 3.2. Effects of urban agriculture and wastewater irrigation on soil microorganisms

#### 3.2.1. Microbial diversity and functionality

DNA sequence read metrics of the six metagenomic samples are reported in Table S1. The taxonomic analysis of the microbial communities at kingdom level in irrigated and non-irrigated fields showed that highest proportion of metagenomic reads mapped to Bacteria (99.1% and 99.2%), followed by Archaea (0.48% and 0.42%), Eukaryota (0.26% and 0.34%), and unassigned (0.05% and 0.06%). In samples of both farming systems, 0.01% of the metagenomics reads were assigned to viruses. Ten and nine bacterial phyla with a relative prevalence ≥ 0.5% of the reads were identified in irrigated and non-irrigated fields, respectively (Figure 1a). The dominant bacteria phyla were in both farming systems *Proteobacteria, Actinobacteria* and *Bacteroidetes* (≥ 89.6% of all bacterial phyla). Phyla abundance was different and not in the same order in the different blocks. Relative prevalence of *Proteobacteria* and *Bacteroidetes* was greater in irrigated soils. The phylum *Gemmatinodetes* was not among the most prevalent phyla in irrigated fields. The Figure 1b reports the top 22 bacterial families found in the different farming systems. Bacterial family’s relative abundance and order were also somewhat different in the farming systems. The 10 most abundant bacterial families in irrigated fields were 1. *Xanthomonadaceae*, 2. *Caulobacteraceae*, 3. *Comamonadaceae*, 4. *Sphingomonadaceae*, 5. *Flavobacteriaceae*, 6. *Mycobacteriaceae*, 7. *Pseudomonadaceae*, 8. *Planctomycetaceae*, 9. *Bradyrhizobiaceae*, and 10. *Nocardioidaceae*. The top 10 in non-irrigated fields were (deviating numbers from irrigated fields are added in brackets) 1. (2) *Caulobacteraceae*, 2. (1) *Xanthomonadaceae*, 3. *Comamonadaceae*, 4. (7) *Pseudomonadaceae*, 5. *Flavobacteriaceae*, 6. (-) *Conexibacteraceae*, 7. (6) *Mycobacteriaceae*, 8. (9) *Bradyrhizobiaceae*, 9. (8) *Planctomycetaceae and* 10. *Nocardioidaceae.*

**Figure 1.**
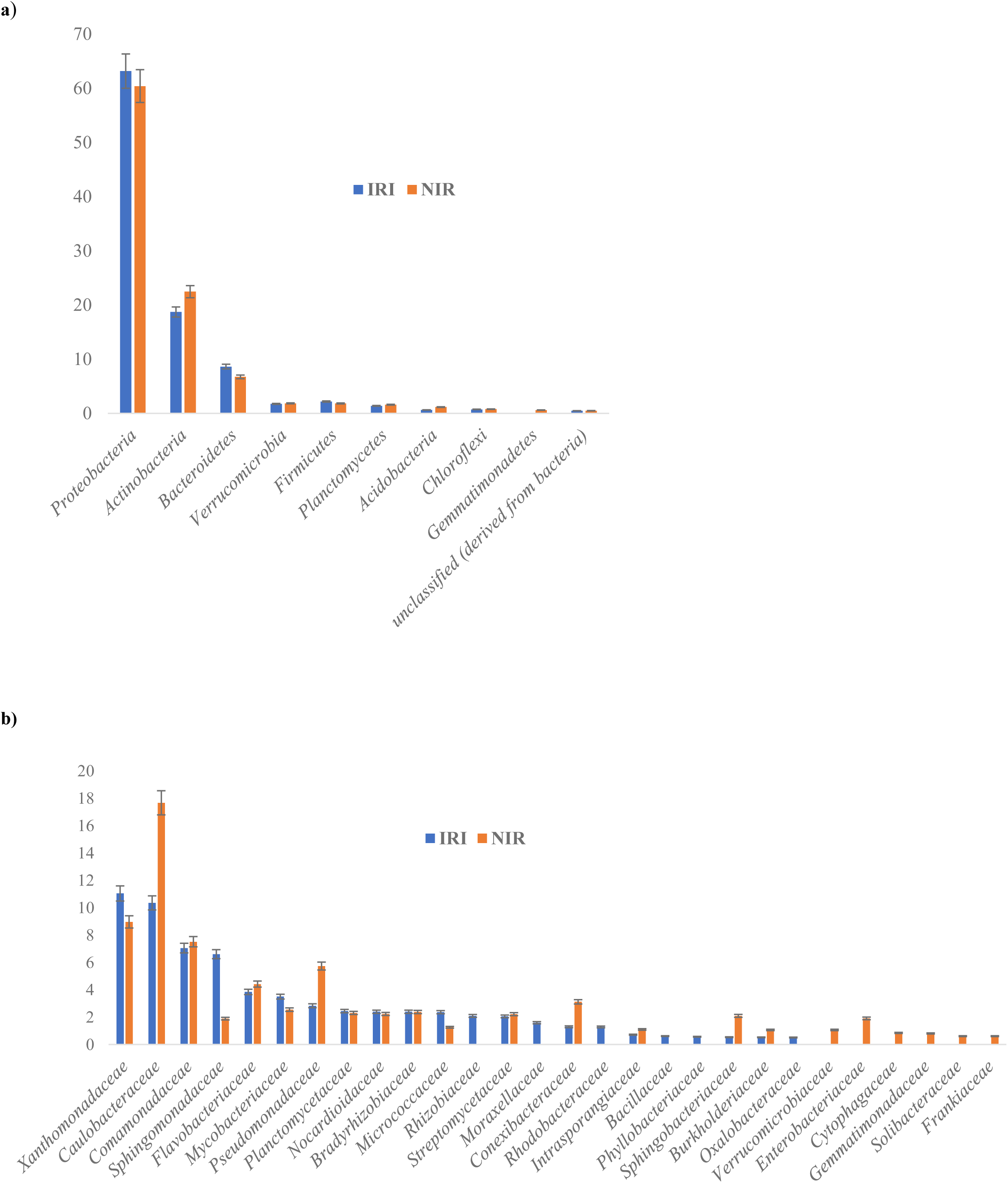
Relative abundances (%) of soil a) bacterial phyla and b) families derived from the metagenomic reads in irrigated fields (IRI) and non-irrigated fields. Bacterial phyla and families with average relative abundance > 0.5% are visualized. (n=3)

Functional metabolic diversity analysis of the six metagenome reads from the soil samples using the SEED database revealed that 14 subsystems were most frequent in the soil microbial communities (Table 2). The most prevalent functional categories were in both farming systems “Carbohydrates”, “Clustering-based subsystems” and “Amino acids and derivatives”. Comparative analysis using the Student’s *t*-test showed that sequence reads coding for functional subsystems ‘’Clustering-based subsystems’’, ‘’DNA metabolism’’, ‘’Nucleosides and nucleotides’’ and ‘’Stress response’’ were significantly higher in irrigated fields (*P* < 0.05).

**Table 2.** Relative distribution of sequencing reads (RPKM units) in major level 1 subsystems in irrigated and non-irrigated fields. Metagenomic data were annotated against SEED subsystems in MG-RAST at a cut off of E-value < 10^-5^.

| Functional subsystems | Agricultural fields |  |  |  |  |  |  |  |
| --- | --- | --- | --- | --- | --- | --- | --- | --- |
|  | Irrigated |  |  |  | Non-irrigated |  |  |  |
|  | IRI1 | IRI2 | IRI3 | Mean (±SD) | NIR1 | NIR2 | NIR3 | Mean (±SD) |
| Carbohydrates | 2.62x 10<sup>6</sup> | 2.35x 10<sup>6</sup> | 2.35x 10<sup>6</sup> | 2.44x 10<sup>6</sup> (±1.56x 10<sup>5</sup>) | 2.51x 10<sup>6</sup> | 2.11x 10<sup>6</sup> | 2.59x 10<sup>6</sup> | 2.40x 10<sup>6</sup> (±2.57x10<sup>5</sup>) |
| Clustering-based subsystems* | 2.55x 10<sup>6</sup> | 2.46x 10<sup>6</sup> | 2.48x 10<sup>6</sup> | 2.50x 10<sup>6</sup> (±4.59x 10<sup>4</sup>) | 2.63x 10<sup>6</sup> | 1.90x 10<sup>6</sup> | 2.82x 10<sup>6</sup> | 2.45x 10<sup>6</sup> (±4.88x 10<sup>5</sup>) |
| Amino acids and derivatives | 2.12x 10<sup>6</sup> | 2.08x 10<sup>6</sup> | 2.08x 10<sup>6</sup> | 2.09x 10<sup>6</sup> (±2.49x 10<sup>4</sup>) | 2.14x 10<sup>6</sup> | 1.65x 10<sup>6</sup> | 2.38x 10<sup>6</sup> | 2.06x 10<sup>6</sup> (±3.74x 10<sup>5</sup>) |
| Miscellaneous | 1.34x 10<sup>6</sup> | 1.33x 10<sup>6</sup> | 1.38x 10<sup>6</sup> | 1.35x 10<sup>6</sup> (±2.58x 10<sup>4</sup>) | 1.41x 10<sup>6</sup> | 9.85x 10<sup>5</sup> | 1.61x 10<sup>6</sup> | 1.33x 10<sup>6</sup> (±3.19x 10<sup>5</sup>) |
| Protein metabolism | 1.31x 10<sup>6</sup> | 1.21x 10<sup>6</sup> | 1.23x 10<sup>6</sup> | 1.25x 10<sup>6</sup> (±5.10x 10<sup>4</sup>) | 1.32x 10<sup>6</sup> | 9.68x 10<sup>5</sup> | 1.40x 10<sup>6</sup> | 1.23x 10<sup>6</sup> (±2.27x 10<sup>5</sup>) |
| Cofactors, vitamins, prosthetic groups, pigments | 1.24x 10<sup>6</sup> | 1.19x 10<sup>6</sup> | 1.20x 10<sup>6</sup> | 1.21x 10<sup>6</sup> (±2.33x 10<sup>4</sup>) | 1.25x 10<sup>6</sup> | 9.46x 10<sup>5</sup> | 1.39x 10<sup>6</sup> | 1.20x 10<sup>6</sup> (±2.27x 10<sup>5</sup>) |
| RNA metabolism | 8.85x 10<sup>5</sup> | 9.30x 10<sup>5</sup> | 9.76x 10<sup>5</sup> | 9.30x 10<sup>5</sup> (±4.58x 10<sup>4</sup>) | 9.84x 10<sup>5</sup> | 6.15x 10<sup>5</sup> | 1.14x 10<sup>6</sup> | 9.13x 10<sup>5</sup> (±2.70x 10<sup>5</sup>) |
| Fatty acids, lipids and isoprenoids | 8.01x 10<sup>5</sup> | 8.19x 10<sup>5</sup> | 7.60x 10<sup>5</sup> | 7.93x 10<sup>5</sup> (±3.01x 10<sup>4</sup>) | 8.26x 10<sup>5</sup> | 6.38x 10<sup>5</sup> | 8.54x 10<sup>5</sup> | 7.73x 10<sup>5</sup> (±1.17x 10<sup>5</sup>) |
| DNA metabolism* | 7.20x 10<sup>5</sup> | 7.23x 10<sup>5</sup> | 7.39x 10<sup>5</sup> | 7.27x 10<sup>5</sup> (±1x 10<sup>4</sup>) | 7.60x 10<sup>5</sup> | 5.06x 10<sup>5</sup> | 8.48x 10<sup>5</sup> | 7.05x 10<sup>5</sup> (±1.78x 10<sup>5</sup>) |
| Cell wall and capsule | 6.69x 10<sup>5</sup> | 7.23x 10<sup>5</sup> | 7.61x 10<sup>5</sup> | 7.18x 10<sup>5</sup> (±4.62x 10<sup>4</sup>) | 7.38x 10<sup>5</sup> | 4.86x 10<sup>5</sup> | 8.76x 10<sup>5</sup> | 7.00x 10<sup>5</sup> (±1.98x 10<sup>5</sup>) |
| Respiration | 4.84x 10<sup>5</sup> | 4.42x 10<sup>5</sup> | 4.34x 10<sup>5</sup> | 4.53x 10<sup>5</sup> (±2.72x 10<sup>4</sup>) | 4.70x 10<sup>5</sup> | 3.81x 10<sup>5</sup> | 4.94x 10<sup>5</sup> | 4.48x 10<sup>5</sup> (±5.93x 10<sup>4</sup>) |
| Nucleosides and nucleotides* | 4.54x 10<sup>5</sup> | 4.37x 10<sup>5</sup> | 4.46x 10<sup>5</sup> | 4.45x 10<sup>5</sup> (±8.73x 10<sup>3</sup>) | 4.63x 10<sup>5</sup> | 3.45x 10<sup>5</sup> | 5.04x 10<sup>5</sup> | 4.37x 10<sup>5</sup> (8.25x 10<sup>4</sup>) |
| Virulence, disease and defense | 4.36x 10<sup>5</sup> | 5.13x 10<sup>5</sup> | 5.85x 10<sup>5</sup> | 5.11x 10<sup>5</sup> (±7.48x 10<sup>4</sup>) | 5.24x 10<sup>5</sup> | 3.12x 10<sup>5</sup> | 7.46x 10<sup>5</sup> | 5.27x 10<sup>5</sup> (±2.17x 10<sup>5</sup>) |
| Stress response* | 4.06x 10<sup>5</sup> | 4.20x 10<sup>5</sup> | 4.23x 10<sup>5</sup> | 4.16x 10<sup>5</sup> (±9.04x 10<sup>3</sup>) | 4.44x 10<sup>5</sup> | 2.93x 10<sup>5</sup> | 4.99x 10<sup>5</sup> | 4.12x 10<sup>5</sup> (±1.07x 10<sup>5</sup>) |

\* The distribution is significantly different between the two agricultural systems (Student's *t*-test, *P* < 0.05). *SD* Standard Deviation. *IRI* Irrigated; *NIR* Non-irrigated.

#### 3.2.2. Antibiotic resistance genes and plasmid replicons

ARGs commonly associated with mobile genetic elements accounted for 33 and 26 out of the 45 and 39 detected ARGs in irrigated and non-irrigated fields sequence reads, respectively (Table 3). The transferable ARGs confer resistance to trimethoprim (2) and nine major classes of antibiotics. They encode resistance to aminoglycosides (10), β-lactams (7), amphenicols (6), tetracyclines (5), sulphonamides (3), macrolides (2), quinolones (1), phosphonic antibiotics (1) and nucleoside antibiotics (1). Twenty-one were common to both farming systems, twelve (*aac*(6’)-Ib7, *ant*(9)-Ia, *cat*III, *cat*Q, *^bla^*CARB-3, *^bla^*OXA-347, *^bla^*OXA-5, *^bla^*Rm3, *fos*B, *sul*3, *tet*C and *tet*X) and five (*mph*G, *^bla^*LCR-1, *ere*A2, *qnr*VC1, and *tet*B(P)) were found in irrigated and non-irrigated fields, respectively. The relative prevalence of ARGs common to both farming systems did not show significant difference (*P*<0.05). Bivariate correlation analysis between prevalence of both ARGs and antibiotic residues showed positive correlations between concentration of sulfamethoxazole, ciprofloxacin, enrofloxacin, trimethoprim and some ARGs (Table 4). Concentration of sulfamethoxazole and ciprofloxacin had the greatest number of positive relationships (nine), followed by enrofloxacin (eight), and trimethoprim (one). Trimethoprim was positively correlated to *dfrA1*. Sulfamethoxazole and ciprofloxacin were positively correlated to *catIII, floR, ^bla^*OXA-347, *^bla^*OXA-5, *^bla^*CARB-3, *^bla^*rm3, *sul3, tetC*, and *tetX*; enrofloxacin was positively correlated to the same ARGs, except *floR.*

**Table 3.**
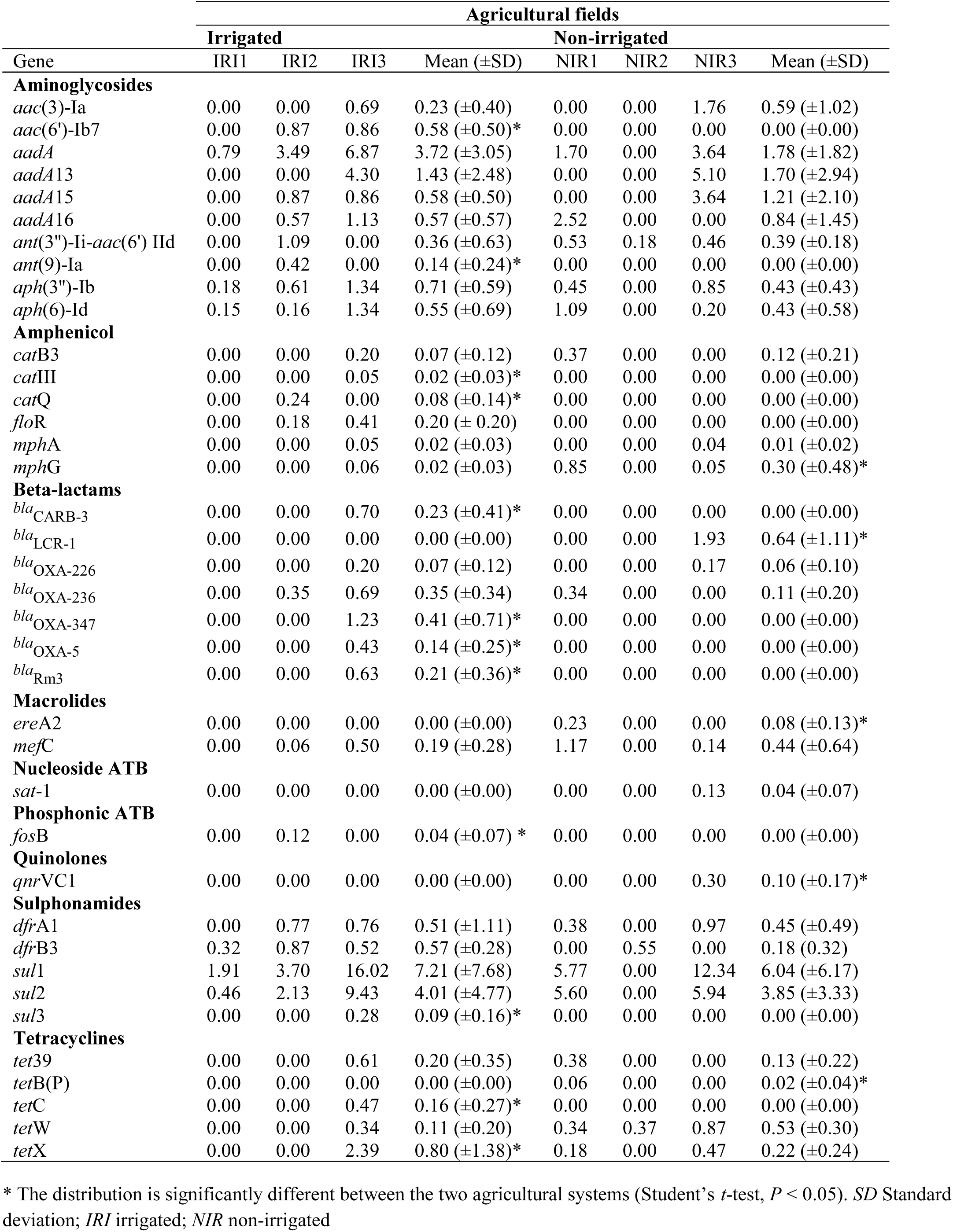
Relative abundance (RPKM units) of the transmissible antibiotic resistance genes and their corresponding families identified in irrigated and non-irrigated fields.

**Table 4.** Positive correlations between relative abundances of antibiotic resistant genes and antibiotics concentrations. In each cell, the value represents Spearman’s coefficient (*r*).

|  | *bla*<sub>CARB-3</sub> | *catIII* | *dfrA1* | *floR* | *bla*<sub>OXA-347</sub> | *bla*<sub>OXA-5</sub> | *bla*<sub>rm3</sub> | *sul3* | *tetC* | *tetX* |
| --- | --- | --- | --- | --- | --- | --- | --- | --- | --- | --- |
| Sulfamethoxazole | 0.979** | 0.979** | - | 0.864* | 0.979** | 0.979** | 0.979** | 0.979** | 0.979** | 0.946** |
| Ciprofloxacin | 0.926** | 0.926** | - | 0.899* | 0.926** | 0.926** | 0.926** | 0.926** | 0.926** | 0.881* |
| Enrofloxacin | 0.970** | 0.970** | - | - | 0.970** | 0.970** | 0.970** | 0.970** | 0.970** | 0.936** |
| Trimethoprim | - | - | 0.869* | - | - | - | - | - | - | - |

\*Asterisk denoted significant correlation at $p < 0.05$ level (2- tailed).
\*\*Double asterisk denoted significant correlation at $p < 0.01$ level (2- tailed).

The mechanisms of resistance encoded by the identified ARGs were dominated by antibiotic inactivation enzymes (64.7% and 71.9%), followed by antibiotic target replacement (14.7% and 12.5%), antibiotic target protection (11.8% and 9.4%) and efflux pumps (6.3% and 8.8%) in irrigated and non-irrigated fields, respectively (Figure S1). ARGs encoding resistance by antibiotic inactivation enzymes was 6% lower in non-irrigated fields, those encoding the other mechanisms of resistance were 2% higher in irrigated fields.

Ten *Enterobacteriaceae* plasmid amplicon incompatibility (Inc) groups were identified in the agricultural fields (Figure 2). Five (IncQ2 ColE IncFIC IncQ1, and IncFII) were common to both, two (IncW IncP1) and three (IncY IncFIB IncFIA) were found in irrigated and non-irrigated fields, respectively.

**Figure 2:**
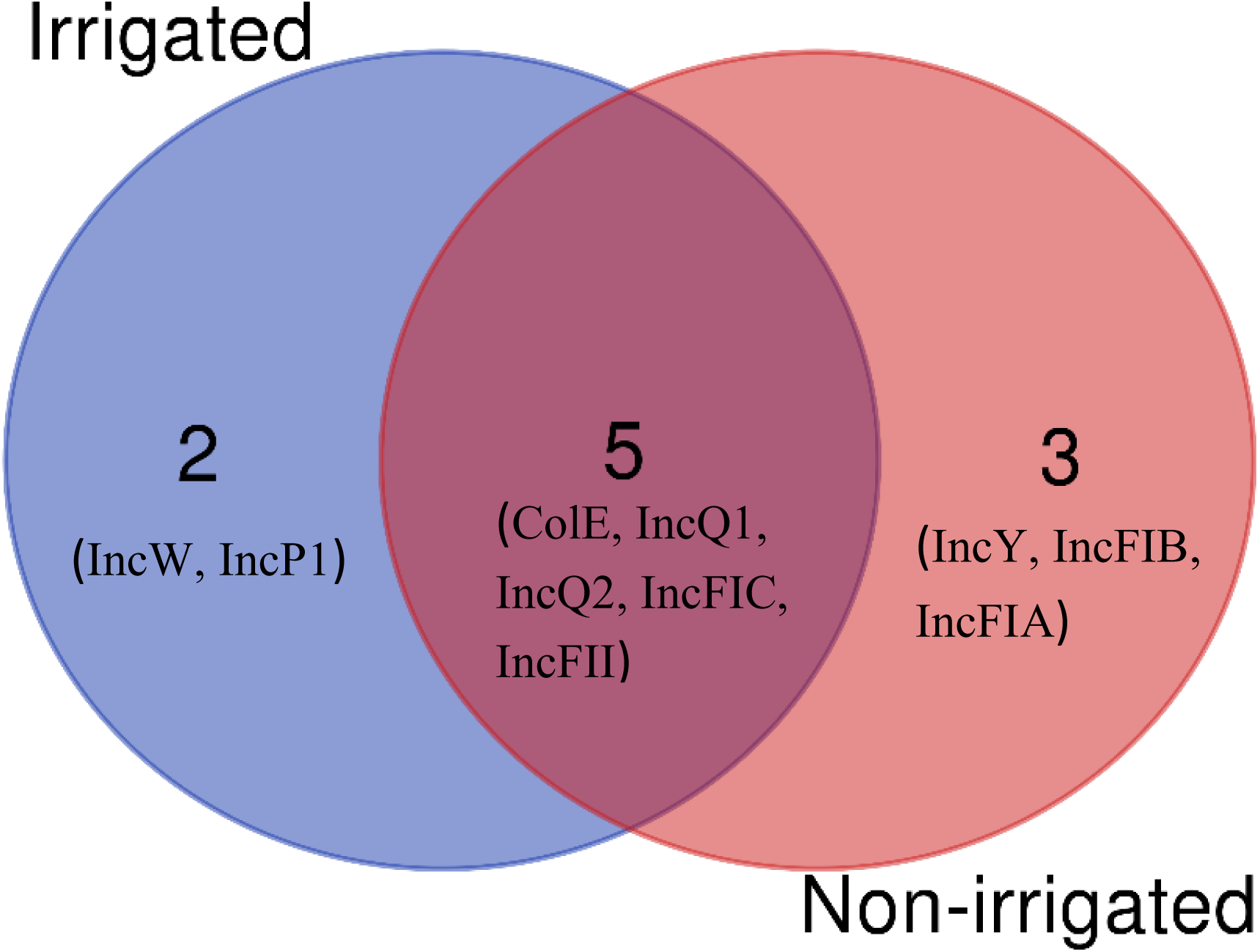
Venn diagram showing the identified *Enterobacteriaceae* plasmid replicons in the irrigated and non-irrigated agricultural fields. (n=3)

## 4. Discussion

The 20 years irrigation of soils with raw wastewater containing substantial amounts of organic matter resulted in higher pH and increased contents of organic carbon and nitrogen compared to non-irrigated soils. The wastewater derived organic matter is subsequently degraded and transformed to soil organic matter (SOM), respectively (Bougnom et al. 2009). Concomitantly increased the residual contents of antibiotics in the irrigated soils. This coincided with previous studies that reported the accumulation of pharmaceuticals in agricultural fields irrigated with treated or raw wastewater (Calderon-Preciado et al., 2011; Grossberger et al. 2014; Wang et al. 2014). The dominant factors influencing the retardation of antibiotics in soil include soil pH, SOM content, and soil texture (Thiele-Bruhn et al., 2004; Du and Liu, 2012). Antibiotic residues are less bioavailable, and thus less biodegradable in soils with high SOM and clay content, owing to stronger sorption to SOM and the formation of non-extractable residues (Luo et al., 2011; Müller et al., 2013; Cheng et al., 2016). Both higher SOM content and higher contents of enrofloxacin, oxytetracycline and sulfamethoxazole residues coincided in irrigated fields. The prevalence of fluoroquinolones and of sulfamethoxazole likely depends on the input but may be also related to the stronger retardation of these antibiotics in acidic and iron oxide-rich tropical soils (Essington et al. 2010) as were investigated in this study. Antibiotic residues were also determined in non-irrigated farms but at lower contents, though. This contamination could be due to previous (not reported to us) use of animal manure as fertilizer, deposition of wastewater aerosol and soil dust, respectively, derived from nearest by irrigated sites by wind erosion, and human transport of agricultural materials between fields (Dalkmann et al. 2012).

Soil microbial communities’ diversity and functions are influenced by abiotic and biotic factors, such as soil texture, pH, carbon content, nutrients, pollutants, and agricultural management (Jangid et al. 2011; Kuramae et al. 2011). In this context it is noted that irrigation wastewater contains nutrient elements and degradable organic material as well as chemical, physical and biological pollutants originating from human activities (Deblonde et al. 2011). Since many pollutants have stimulating or inhibiting effect on microbial cells, it was not surprising to find changes in both soil microbial composition and function following irrigation with wastewater.

The identified bacterial phyla in both irrigated and non-irrigated fields are generally encountered in soil (Fierer et al. 2012; Nacke et al. 2014). Previous field and mesocosm-scale studies have shown the transfer of *Bacteroidetes* and *Proteobacteria* phyla members into the soil following wastewater irrigation (Broszat et al., 2014; Frenk et al., 2018). The increased number of these phyla is explained by their copiotrophic activity and high grow rates in the presence of nutrients, moisture and labile organic C found in wastewater (Broszat et al. 2014). This elucidates their greater prevalence in irrigated fields. *Acidobacteria, Chloroflexi, Verrucomicrobia* and *Planctomycetes* have been reported to follow oligotrophic strategies with limited growth rates and thriving capacity in nutrient-poor ecosystems (Lauber et al. 2013; Kielak et al. 2016). The members of *Xanthomonadaceae* family exhibit higher abundance in wastewater irrigated fields, and those of the *Caulobacteraceae* family are primarily known as oligotrophic microorganisms (Poindexter et al., 1981). Tropical soils have high mineralisation rate and are poor in nutrients. This could explain why these bacterial phyla and families are prevalent in non-irrigated fields.

The analysis of metagenome reads using SEED provided insights into the functional metagenomic profiling of microorganisms living in the investigated fields. Considering changes in the microbial diversity structure observed between the two farming systems, some differences in the metabolic potential of the soil microbiota were expected. The functions ‘clustering-based subsystems’ (functional coupling evidence but unknown function), ‘DNA metabolism’ (DNA repair, bacterial), ‘nucleosides and nucleotides and stress response’ translate a higher bacterial and enzymatic activity in irrigated fields owing to the introduction of wastewater that is rich in nutrients, organic matter, and contains several pollutants. The soil microbiota must develop functional redundancy and adopt mechanisms to adapt, survive and grow. Therefore, wastewater irrigation affects microbial community structure and functions.

Antibiotic resistance is an ancient phenomenon and soil is the potentially largest reservoir of genes coding for antibiotic resistance owing to its complex microbial community able to produce antibiotic compounds (Nesme and Simonet, 2015). Relevant quantity and diversity of the genes coding for bacterial resistance have been already uncovered from soils (Heuer et al., 2011, Dalkmann et al. 2012). The determination of transferable ARGs in the investigated soils is in accordance with previous studies (Nesme et al. 2014; D’Costa et al. 2011). Furthermore, an additional effect of wastewater irrigation on the soil resistome was found. Studies of Chen et al. (2016); Broszat et al. (2014); Dungan et al. 2018 in China, Mexico, in US, have reported that wastewater irrigation enhanced the occurrence of ARGs in soils. Some pathways likely to induce bacterial resistance in soil include, influx of antibiotic residues which can induce a selective pressure; transfer and survival of ARB; and influx of transferable plasmids harbouring ARGs (Von Wintersdorff et al. 2016). The wastewaters used to irrigate the investigated fields were previously reported to be strong vector of bacterial resistance (Bougnom et al. 2019a; 2019b); thus, a more diverse ARB community and ARGs in irrigated fields was anticipated. All the ARGs present in the fields were already reported in the wastewaters. However, the diversity of ARGs found in the soil was lower. This is most likely a consequence of the death of many ARB when transferred from wastewater to soil where they must develop mechanisms to adapt, survive and growth. The surviving bacterial resistome can disseminate drug-resistance in the environment. Among the ARGs found in irrigated fields, four out of 12 were genes encoding extended spectrum β-lactamase. Extended-spectrum β-lactamases (ESBLs) are primarily carried by Gram-negative bacteria, and manure application to soil sustains the survival and growth of pathogens, ARB and ARGs (Rawat and Nair, 2010; Sharma and Reynnells, 2016). *Enterobacteriaceae* producing ESBLs can therefore live for longer times in irrigated fields and contaminate humans and animals via direct contact or the food chain. ESBLs producers pose critical issues on clinical settings since they are able to inactivate β-lactams, thus requiring the administration of more expensive antibiotics. Urban farmworkers and consumers feeding with crops coming from these fields might be exposed to serious health risks. The modification of the different percentages of antibiotic resistance mechanisms in the irrigated field was consequent to the modification of the soil resistome following wastewater irrigation. Soils are important reservoirs of diverse ARGs and application of manure, sludge and irrigation by groundwater could introduce ARB and ARGs in soils (Binh et al. 2008; Negreanu et al. 2012; Nesme and Simonet, 2015). This could explain the presence of transferable ARGs in non-irrigated fields. The relative prevalence of *Enterobacteriaceae* family was greater in non-irrigated farms, which suggests the use of animal manure for soil fertilization.

Concentrations of sulfamethoxazole and trimethoprim were positively correlated to relative prevalence of *sul3* and *dfrA1*, respectively, suggesting their influence on the selection of these genes. Concentration of antibiotics residue in the environment exerting a selective pressure on the acquisition of ARGs coding resistance against them are in accordance with studies of Cheng et al. (2016) and Pan and Chu (2018). We found that sulfamethoxazole, ciprofloxacin and enrofloxacin selected for multiple and cross resistance. Their presence at a critical level could foster the maintenance of foreign resistance genes not oriented against their action and/or select for multidrug resistance plasmids (Levy, 2002; Blanco et al., 2016). Studies by Botts et al. (2017) in the US have reported multidrug resistant plasmids encoding resistance genes for tetracyclines, sulfonamides, β-lactams, fluoroquinolones, aminoglycosides, and amphenicols in coastal wetland impacted by urban stormwater runoff and wastewater. Therefore, if the concentration of one of these antibiotics is high enough in the environment, it can exert a selective pressure on plasmid carrying genes conferring resistance to those antibiotics. This shows the complex nature of factors selecting ARB and ARGs in the environment.

Considering the wide and diverse ARGs in the wastewaters used to irrigate these fields (Bougnom et al. 2019b), *Enterobacteriaceae* plasmid replicons were expected. However, the different amplicon groups found in the fields were less diverse that the ones found in the wastewater. This confirms the death of many ARB following their transfer from wastewater to soil or loss of some ARGs considering the fitness cost for their maintenance. IncFIB has been reported to be predominant in cattle faeces in Nigeria (Inwezerua et al. 2014); thus, fertilisation with animal manure could explain plasmid replicons in non-irrigated fields. Considering the epidemiological aspects, *Enterobacteriaceae* plasmids replicons found in irrigated farms represent a greater public health issue. IncP and IncW type plasmids can transfer and maintain themselves in all Gram-negative bacteria (Dröge et al., 2000), while IncF and incY have a “narrow” host range (Boyd et al. 1996; Mshana et al. 2009).

## Conclusion

This is the first study using metagenomics to investigate the impact of raw wastewater use for urban agriculture on bacterial resistance in soil on example of sites in three major cities in Africa. Our data revealed that raw wastewater irrigation impacts both soil microbial communities and functional abilities. Microbial communities living in irrigated fields must develop mechanisms to adapt, survive and grow. Critical concentrations of sulfamethoxazole, ciprofloxacin and enrofloxacin antibiotic residues may select for ARGs not oriented against their action. Wastewater irrigation favours the presence of more diverse ARGs of clinical relevance in soil. The diversity of ARGs present in irrigated fields is lower than the one found in the wastewaters. *Enterobacteriaceae* plasmid replicon groups found in irrigated fields present a significant public health concern. The collected information suggests that raw wastewater irrigated soils in Africa could represent a vector for the spread of antibiotic resistance, thus, threatening human and animal health. Additional studies are needed to investigate the consequence of exposure for farmworkers and food consumers.

## Acknowledgments

We thank the European Commission; this project has received funding from the European Union’s Horizon 2020 research and innovation programme under the Marie Sklodowska-Curie grant agreement No 655398. We are grateful to Petra Ziegler and Elvira Sieberger, University of Trier, for assistance in the physical and chemical analyses, and members of the Antimicrobials Research Group, University of Birmingham, for helpful and constructive discussions.

## References

Adegoke, A.A., Isaac D. Amoah, I.D., Thor A. Stenström, T.A., Verbyla, M.E., Mihelcic, J.R., 2018. Epidemiological evidence and health risks associated with agricultural reuse of partially treated and untreated wastewater: A review. Front Public Health. 6, 337.

Alves, L.D.F., Westmann, C.A., Lovate, G.L., de Siqueira, G.M.V., Borelli, T.C., Guazzaroni, M., 2018. Metagenomic approaches for understanding new concepts in microbial science. Int. J. Genomics. 2312987.

Amos, G.C.A., Hawkey, P.M., Gaze, W.H., Wellington, E.M., 2014. Waste water effluent contributes to the dissemination of CTX-M-15 in the natural environment. J. Antimicrob. Chemoth. 69, 1785–1791.

Andersson, D.I., Hughes, D., 2010. Antibiotic resistance and its cost: is it possible to reverse resistance? Nat. Rev. Microbiol. 8, 260–271.

Asnicar, F., Weingart, G., Tickle, T.L., Huttenhower, C., Segata, N., 2015. Compact graphical representation of phylogenetic data and metadata with GraPhlAn. Peerj 3.

Blackwell, P.A., Holten Lutzhoft, H.C., Ma, H.P., Halling-Sørensen, B., Boxall, A.B.A., Kay, P., 2004. Ultrasonic extraction of veterinary antibiotics from soils and pig slurry with SPE clean-up and LC-UV and fluorescence detection. Talanta 64,1058-64.

Blanco, P., Hernando-Amado, S., Reales-Calderon, J., Corona, F., Lira, F., Alcalde-Rico, M., Bernardini, A,, Sanchez, M.B., Martinez, J.L., 2016. Bacterial multidrug efflux pumps: much more than antibiotic resistance determinants. Microorganisms 4:E14.

Binh, C.T.T., Heuer, H., Kaupenjohann, M., Smalla. K., 2008. Pig-gery manure used for soil fertilization is a reservoir for transferable antibiotic resistance plasmids. FEMS Microbiol. Ecol. 66:25–37.

Botts, R.T., Apffel, B.A., Walters, C.J., Davidson, K.E., Echols, R.S., Geiger, M.R., Guzman, V.L., Haase, V.S., Montana, M.A., La Chat, C.A., Mielke, J.A., Mullen, K.L., Virtue, C.C, Brown, C.J., Top, E.M., Cummings, D.E., 2017. Characterization of four multidrug resistance plasmids captured from the sediments of an urban coastal wetland. Front. Microbiol. 8:1922.

Bougnom, B. P., Zongo C., McNally A., Ricci V., Etoa F.X., Thiele-Bruhn S., Piddock L.J.V., 2019a. Wastewater used for urban agriculture in West Africa as a reservoir for antibacterial resistance dissemination. Environ Res 168, 14–24.

Bougnom, B.P., McNally, A., Etoa F.X., PiddocK, L.J.V., 2019b. Urban population density impacts on the abundance of antibiotic resistance genes, except those encoding ESBL in raw sewage used for urban agriculture in Africa (Accepted in Environmental Pollution)

Boyd, E.F., Hill, C.W., Rich, S.M., Hartl, D.L., 1996. Mosaicstructure of plasmids from natural populations of *Escherichia coli*. Genetics 143, 1091–1100.

Broszat, M., Nacke, H., Blasi, R., Siebe, C., Huebner, J., Daniel, R., Grohmann, E., 2014. Wastewater irrigation increases abundance of potentially harmful Gammaproteobacteria in soils from Mezquital Valley, Mexico. Appl. Environ. Microbiol. 80, 5282–5291.

Buchholz, U., Bernard, H., Werber, D., Böhmer, M.M., Remschmidt, C.,Wilking, H., Deleré, Y., an der Heiden, M., Adlhoch, C., Dreesman, J., Ehlers, J., Ethelberg, S., Faber, M., Frank, C, Fricke, G, Greiner, M, Höhle, M, Ivarsson, S, Jark, U, Kirchner, M, Koch, J, Krause, G., Luber, P, Rosner, B., Stark, K., Kühne, M. 2011. German outbreak of *Escherichia coli* O104:H4associated with sprouts. N. Engl. J. Med. 365, 1763–1770.

Calderón-Preciado, D., Jiménez-Cartagena, C., Matamoros, V., Bayona, J.M., 2011. Screening of 47 organic microcontaminants in agricultural irrigation waters and their soil loading. Water Res. 45, 221–231.

Carattoli, A., Zankari, E., Garcia-Fernandez, A., Larsen, M.V., Lund, O., Villa, L., Aarestrup, F.M. and Hasman, H., 2014. In Silico Detection and Typing of Plasmids using PlasmidFinder and Plasmid Multilocus Sequence Typing. Antimicrob Agents Ch 58, 3895–3903.

Cheng, W.X., Li, J.N., Wu, Y., Lk, Xu, Su, C., Qian, Y.Y., et al., 2016. Behavior of antibiotics and antibiotic resistance genes in eco-agricultural system: a case study. J. Hazard. Mater. 304, 18–25.

Dalkmann, P., Broszat, M., Siebe, C., Willaschek, E., Sakinc, T., Huebner, J., Amelung, W., Grohmann, E., Siemens, J., 2012. Accumulation of pharmaceuticals, *enterococcus*, and resistance genes in soils irrigated with wastewater for zero to 100 years in central Mexico. PLoS One 7 (9), e45397.

D’Costa, V.M., King, C.E., Kalan, L., Morar, M., Sung, W.W., Schwarz, C., Froese, D., Zazula, G., Calmels, F., Debruyne, R., Golding, G.B., Poinar, H.N., Wright, G.D., 2011. Antibiotic resistance is ancient. Nature 477, 457–461.

Deblonde, T., Cossu-Leguille, C., Hartemann, P., 2011. Emergingpollutants in wastewater: a review of the literature. Int. J. Hyg. Environ. Heal. 214, 442–448.

Dickin, S.K., Schuster-Wallace, C.J., Qadir, M., Pizzacalla, K., 2016. A Review of Health Risks and Pathways for Exposure to Wastewater Use in Agriculture. Environ. Health. Persp. 124, 900–909.

Droge, M., Puhler, A., Selbitschka, W., 2000. Phenotypic and molecular characterization of conjugative antibiotic resistance plasmids isolated from bacterial communities of activated sludge. Mol. Gen. Genet. 263, 471–82.

Du, L., Liu, W., 2012. Occurrence, fate, and ecotoxicity of antibiotics in agro-ecosystems. A review. Agron Sustain Dev 32, 309–27.

Dungan, R.S., McKinney, C.W., Leytem, A.B., 2018. Tracking antibiotic resistance genes in soil irrigated with dairy wastewater. Sci. Total Environ. 635, 1477–1483.

Essington, M.E., Lee, J., Seo, Y., 2010. Adsorption of antibiotics by montmorillonite and kaolinite. Soil Sci. Soc. Am. J. 74, 1557–1588.

Fierer, N, Leff, J.W., Adams, B.J., Nielsen, U.N., Bates, S.T., Lauber, C.L., Owens, S., Gilbert, J.A., Wall, D.H., Caporaso, J.G., 2012. Cross-biome metagenomic analyses of soil microbial communities and their functional attributes. Proc. Natl. Acad. Sci. U.S.A. 109, 21390–21395.

Forsberg, K.J., Reyes, A., Bin, W., Selleck, E.M., Sommer, M.O.A., Dantas, G., 2012. The Shared Antibiotic Resistome of Soil Bacteria and Human Pathogens. Science 337, 1107–1111.

Glass E.M., Wilkening J., Wilke, A., Antonopoulos, D., Meyer, F., 2010. Using the metagenomics RAST server (MG-RAST) for analyzing shotgun metagenomes. Cold Spring Harb Protoc: prot 5368.

Grossberger, A., Hadar, Y., Borch, T., Chefetz, B., 2014. Biodegradability of pharmaceutical compounds in agricultural soils irrigated with treated wastewater. Environ. Pollut. 185:168–177.

Heuer H., Schmitt H., Smalla K. 2011. Antibiotic resistance gene spread due to manure application on agricultural fields. Curr. Opin. Microbiol. 14, 236–24310.

Inwezerua, C., Mendonc, N., Calhau, V., Domingues, S., Adeleke, O.E., Da Silva, G.J. 2014. Occurrence of extended-spectrum beta-lactamases in human and bovine isolates of *Escherichia coli* from Oyo state, Nigeria, J. Infect Dev Countr, 8, 774–779.

Jangid, K., Williams, M.A., Franzluebbers, A.J., Sanderlin, J.S., Reeves, J.H., Jenkins, M.B., Endale, D.M., Coleman, D.C., Whitman, W.B., 2008. Relative impacts of land-use, management intensity and fertilization upon soil microbial community structure in agricultural systems. Soil Biology and Biochemistry 40, 2843–2853.

Kaminski, J., Gibson, M.K., Franzosa, E.A., Segata, N., Dantas, G., Huttenhower, C., 2015. High-Specificity Targeted Functional Profiling in Microbial Communities with ShortBRED. Plos Comput Biol 11(12).

Kielak, A., Barreto, C., Kowalchuk, G.A., van Veen J.A., Kuramae, E.E., 2016. The ecology of Acidobacteria: moving beyond genes and genomes. Front Microbiol 7, 00744.

Kuramae, E.E., Yergeau E., Wong, L.C., Pijl, A.S., van Veen, J.A., Kowalchuk, G.A., 2011. Soil characteristics more strongly influence soil bacterial communities than land-use type. FEMS Microbiol. Ecol. 79, 12–24.

Lauber, C.L., Ramirez, K.S., Aanderud, Z., Lennon, J., Fierer, N., 2013. Temporal variability in soil microbial communities across land-use types. ISME J 7, 1641–1650.

Levy, S.B. 2002. Factors impacting on the problem of antibiotic resistance. Journal of Antimicrobial Chemotherapy 49, 25–30.

Luo, Y., Xu, L., Rysz, M., Wang, Y.Q., Zhang, H., Alvarez, P.J.J., 2011. Occurrence and transport of tetracycline, sulfonamide, quinolone, and macrolide antibiotics in the Haihe River basin, China. Environ. Sci. Technol. 45, 1827–1833.

Mateo-Sagasta, J., Medlicott, K., Qadir, M., Rashid-Sally, L., Dreschel, P., Liebe, J., 2013. Proceedings of the UN-Water project on the Safe Use of Wastewater in Agriculture. UNW-DPC Proceedings Series, August 2013. http://collections.unu.edu/eserv/UNU:2661/proceedings-no-11_WEB.pdf.

McArthur, A.G., Waglechner, N., Nizam, F., Yan, A., Azad, M.A., Baylay, A.J., Bhullar, K., Canova, M.J., De Pascale, G., Ejim, L., Kalan, L., King, A.M., Koteva, K., Morar, M., Mulvey, M.R., O’Brien, J.S., Pawlowski, A.C., Piddock, L.J.V., Spanogiannopoulos, P., Sutherland, A.D., Tang, I., Taylor, P.L., Thaker, M., Wang, W.L., Yan, M., Yu, T. and Wright, G.D., 2013. The Comprehensive Antibiotic Resistance Database. Antimicrob Agents Ch 57, 3348–3357.

Michelini L., Reichel R., Werner W., Ghisi R., Thiele-Bruhn S. (2012) Sulfadiazine uptake and effects on Salix fragilis L. and Zea maize L. plants. Water, Air and Soil Pollution 223, 5243–5257.

Mshana, S.E., Imirzalioglu, C., Hossain, H., Hain, T., Domann, E., Chakraborty, T. 2009. Conjugative IncFI plasmids carrying CTX-M-15 among *Escherichia coli* ESBL producing isolates at a University hospital in Germany. BMC Infect Dis 9, 97.

Müller, P., Alber, D.G., Turnbull, L., Schlothauer R.C., Carter D.A., Whitchurch C.B., Harry, E.J., 2013. Synergism between Medihoney and rifampicin against methicillin-resistant *Staphylococcus aureus* (MRSA). PLoS ONE 8, e57679

Nacke, H., Fischer, C., Thürmer, A., Meinicke, P., Daniel, R., 2014. Land use type significantly affects microbial gene transcription in soil. Microb. Ecol. 67:919–930.

Nesme, J., Cécillon, S., Delmont, T.O., Monier, J.M., Vogel, T.M., Simonet. P., 2014. Large-scale metagenomic-based study of antibiotic resistance in the environment. Curr. Biol. 24, 1096–1100.

Nesme, J., Simonet. P., 2015. The soil resistome: A critical review on antibi-otic resistance origins, ecology and dissemination potential in telluric bac-teria. Environ. Microbiol. 17, 913–930.

Pan M., Chu, L.M., 2018. Occurrence of antibiotics and antibiotic resistance genes in soils from wastewater irrigation areas in the Pearl River Delta region, southern China. Sci. Total Environ 624, 145–152.

Piddock LJ., 2012. The crisis of no new antibiotics—what is the way forward? Lancet Infect Dis. 12, 249–253.

Poindexter, J.S., 1981. The Caulobacters: Ubiquitous Unusual Bacteria. Microbiol Rev, 45: 123–179.

Rawat, D., Nair, D., 2010. Extended-spectrum ß-lactamases in Gram negative bacteria. J. Glob. Infect. Dis. 2, 263–74.

Sentchilo, V., Mayer, A.P., Guy, L., Miyazaki, R., Tringe, S.G., Barry, K., Malfatti, S., Goessmann, A., Robinson-Rechavi, M., van der Meer, J.R., 2013. Communitywide plasmid gene mobilization and selection. ISME J. 7, 1173–1186.

Sharma, M., Reynnells, R., 2016. Importance of Soil Amendments: Survival of Bacterial Pathogens in Manure and Compost Used as Organic Fertilizers. Microbiol. Spectrum 4, PFS-0010-2015

Thiele-Bruhn, S., Seibicke, T., Schulten, H.-R., Leinweber, P., 2004. Sorption of sulfonamide pharmaceutical antibioticson whole soils and particle-size fractions. J. Environ. Qual. 33, 1331–1342.

Von Wintersdorff, C.J.H., Penders, J., van Niekerk, J.M., Mills, N.D., Majumder, S., van Alphen, L.B., Savelkoul, P.H.M., Wolffs, P.F.G., 2016. Dissemination of Antimicrobial Resistance in Microbial Ecosystems through Horizontal Gene Transfer. Front. Microbiol. 7.

Wang, F., Qiao, M., Su, J.Q., Chen, Z., Zhou, X., Zhu, Y.G. 2014. High throughput profiling of antibiotic resistance genes in urban park soils with reclaimed water irrigation. Environ. Sci. Technol. 48, 9079–9085.

Wellington, E.M.H., Boxall, A.B.A., Cross, P., Feil, E.J., Gaze, W.H., Hawkey, P.M., Johnson-Rollings, A.S., Jones, D.L., Lee, N.M., Otten, W., Thomas, C.M. and Williams, A.P., 2013. The role of the natural environment in the emergence of antibiotic resistance in Gram-negative bacteria. Lancet Infect Dis 13, 155–165.

