## Supplemental Figure 1 for "Raw wastewater irrigation for urban agriculture in Africa increases the diversity of transferable antibiotic resistances genes in soil, including those encoding ESBLs"

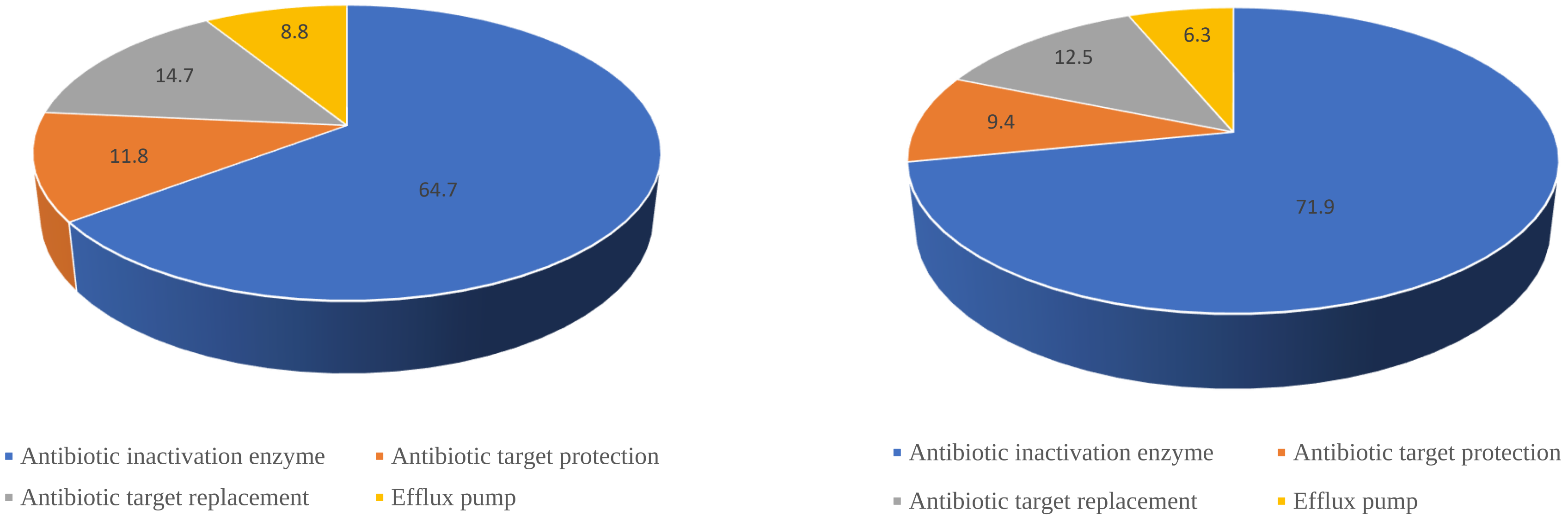


**Figure S1**. Mechanisms of antibiotic resistance (%) of the antibiotic resistance genes (based on their diversity) detected in irrigated and non-irrigated fields (n=3).
