## Supplemental Table 1 for "Raw wastewater irrigation for urban agriculture in Africa increases the diversity of transferable antibiotic resistances genes in soil, including those encoding ESBLs"

**Table S1.** DNA sequence read metrics of the six metagenomic samples from soil from irrigated fields (IRI) and non-irrigated fields (NIR) based on MG-RAST annotation.

| **Metagenome** | | **Sequences count** | **Sequences count post QC** | **Mean GC content post QC** | **Mean sequence length post QC** |
|---|---|---|---|---|---|
| | IRI1 | 3,792,462,000 bp | 3,649,105,747 bp | 65 ± 10 % | 149 ± 3 bp |
| Irrigated | IRI2 | 3,439,886,400 bp | 3,309,468,880 bp | 62 ± 12 % | 149 ± 3 bp |
| | IRI3 | 3,491,007,600 bp | 3,329,257,884 bp | 60 ± 12 % | 149 ± 3 bp |
| | NIR1 | 3,527,491,650 bp | 3,394,411,184 bp | 63 ± 11 % | 149 ± 3 bp |
| Non-irrigated | NIR2 | 3,258,523,350 bp | 3,159,665,932 bp | 66 ± 9 % | 149 ± 3 bp |
| | NIR3 | 4,120,454,250 bp | 3,682,552,830 bp | 62 ± 12 % | 150 ± 3 bp |
